# Metformin Does Not Attenuate Angiotensin II-Induced Aortic Aneurysms in Low-Density Lipoprotein Receptor Deficient Mice

**DOI:** 10.1101/2025.05.25.656029

**Authors:** Samuel C. Tyagi, Sohei Ito, Jacob C. Hubbuch, Michael K. Franklin, Deborah A. Howatt, Hong S. Lu, Alan Daugherty, Hisashi Sawada

## Abstract

**Background:** Metformin, a biguanide antihyperglycemic agent, prevents angiotensin II (AngII)-induced abdominal aortic aneurysm formation in apolipoprotein E-deficient (ApoE−/−) mice. Low-density lipoprotein receptor-deficient (LDLR−/−) mice are also widely used as a hypercholesterolemic model, which more closely mimics the lipoprotein profile of patients with hypercholesterolemia than ApoE−/− mice. In addition, LDLR−/− mice exhibit characteristics of glucose metabolism that are distinct from ApoE−/− mice. However, it remains unknown whether metformin suppresses AngII-induced aortic aneurysm formation in LDLR−/− mice.

**Methods:** Male LDLR−/− mice at 9 weeks of age were administered either vehicle or metformin in drinking water and fed a Western diet. Subsequently, AngII was infused into mice for 4 weeks. Plasma metformin concentrations were measured by mass spectrometry. Maximal aortic diameters and areas were measured ex vivo.

**Results:** Mass spectrometry analysis determined plasma metformin concentrations in mice administered the drug. Metformin administration resulted in lower body weight compared to the vehicle group, indicating effective metformin administration. However, ex vivo measurements demonstrated that metformin did not reduce aortic diameters in the suprarenal abdominal region. In addition, metformin failed to prevent AngII-induced ascending aortic dilatations.

**Conclusion:** Metformin did not attenuate AngII-induced aortic aneurysm formation in either the ascending or suprarenal abdominal region of LDLR−/− mice.

## INTRODUCTION

Abdominal aortic aneurysm (AAA) is a life-threatening vascular disease.[1, 2] The life-threatening sequela of AAA is rupture with the risk of aortic rupture being proportional to the diameter of AAA.[3] The Society for Vascular Surgery guidelines recommend repair when AAA diameter reaches 5.5 cm.[4] Despite technological improvements, open surgery and endovascular procedures remain the only treatment modalities for AAA. There is no approved medical therapy to prevent AAA progression and rupture. In patients with asymptomatic AAA, management primarily focuses on blood pressure control, smoking cessation, and continued surveillance, which highlights the limited progress in developing effective medical therapies for AAA.[4]

A number of both clinical and preclinical studies reported that diabetes promotes many types of vascular diseases, including hypertension and atherosclerosis.[5, 6] However, an inverse relationship exists between diabetes and AAA. Clinical studies have demonstrated consistently that AAA growth is greater in patients without diabetes compared to those with diabetes.[7–9] One possible explanation for the reduced risk of AAA in patients with diabetes is the secondary effects of diabetes medications. Retrospective clinical studies have found that diabetic patients taking metformin, a biguanide antihyperglycemic agent, exhibited slower aneurysm growth and a lower rupture rate compared to those not prescribed metformin.[10–12] Experimentally, metformin protected against AAA formation induced by intra-aortic porcine pancreatic elastase in mice.[10] In addition, metformin prevented angiotensin II (AngII)-induced AAA formation in apolipoprotein E-deficient (ApoE−/−) mice, a hypercholesterolemic mouse model.[13, 14] These findings suggest that metformin may have the potential to suppress AAA formation and progression.

AngII infusion in mice has been used widely to determine mechanisms underlying AAA formation.[15, 16] There is consistent evidence that hypercholesterolemia promotes AngII-induced AAA in both ApoE and LDLR−/− mice.[15, 17] The incidence of AngII-induced AAA is generally less than 20% in normocholesterolemic C57BL/6J mice but exceeds 80% in hypercholesterolemic strains.[16, 18] Of note, ApoE−/− mice and LDLR−/− mice differ in lipoprotein characteristics.[19] LDLR−/− mice develop hypercholesterolemia characterized by increased LDL-cholesterol concentrations, whereas ApoE−/− mice exhibit increased concentrations of VLDL and chylomicron remnants.[19] Therefore, in contrast to ApoE−/− mice, LDLR−/− mice replicate the lipoprotein profile of patients with hypercholesterolemia. In addition, these two strains show differences in glucose metabolism.[20] Considering these lipoprotein and metabolic differences, in this study, we investigated whether metformin suppressed AngII-induced AAA formation in LDLR−/− mice.

## MATERIALS AND METHODS

### Mice

Male LDLR−/− mice were purchased from The Jackson Laboratory (#002207). Because of the low incidence of AngII-induced AAAs in female mice,[16] only male mice were studied. Mice were housed in ventilated cages with negative air pressure (Allentown Inc.). Hardwood chips were used as bedding (TJ Murphy). Drinking water filtered by reverse osmosis and normal mouse laboratory diet (#2918; Harlan Teklad) were provided ad libitum. Rooms were set with a 14/10 hour light/dark cycle, a temperature range of 20-23°C, and a humidity range of 50-60%. Mice were randomly assigned to the study groups, and no mice were excluded from the study. All animal experiments were approved by the University of Kentucky’s Institutional Animal Care (2018-2967) and Use Committee and adhered to the ARRIVE (Animal Research: Reporting of In Vivo Experiments) guidelines.

### Metformin and angiotensin II

The experimental design is illustrated in **Figure 1**. Mice were administered either vehicle or metformin (1.8 mg/mL, Sigma-Aldrich, #D150959-5G) in drinking water starting at 9 weeks of age and continuing throughout the experiment. Three days later, the diet was switched to a Western diet containing milk fat (21% wt/wt) and cholesterol (0.2% wt/wt; TD.88137, Inotiv). After 1week, mini-osmotic pumps (Model 2004, Alzet LLC) were implanted subcutaneously to infuse AngII (1,000 ng/kg/min; #H-1705, Bachem) for 28 days.[15, 21] Mice were randomly assigned to study groups.

**Fig 1.**
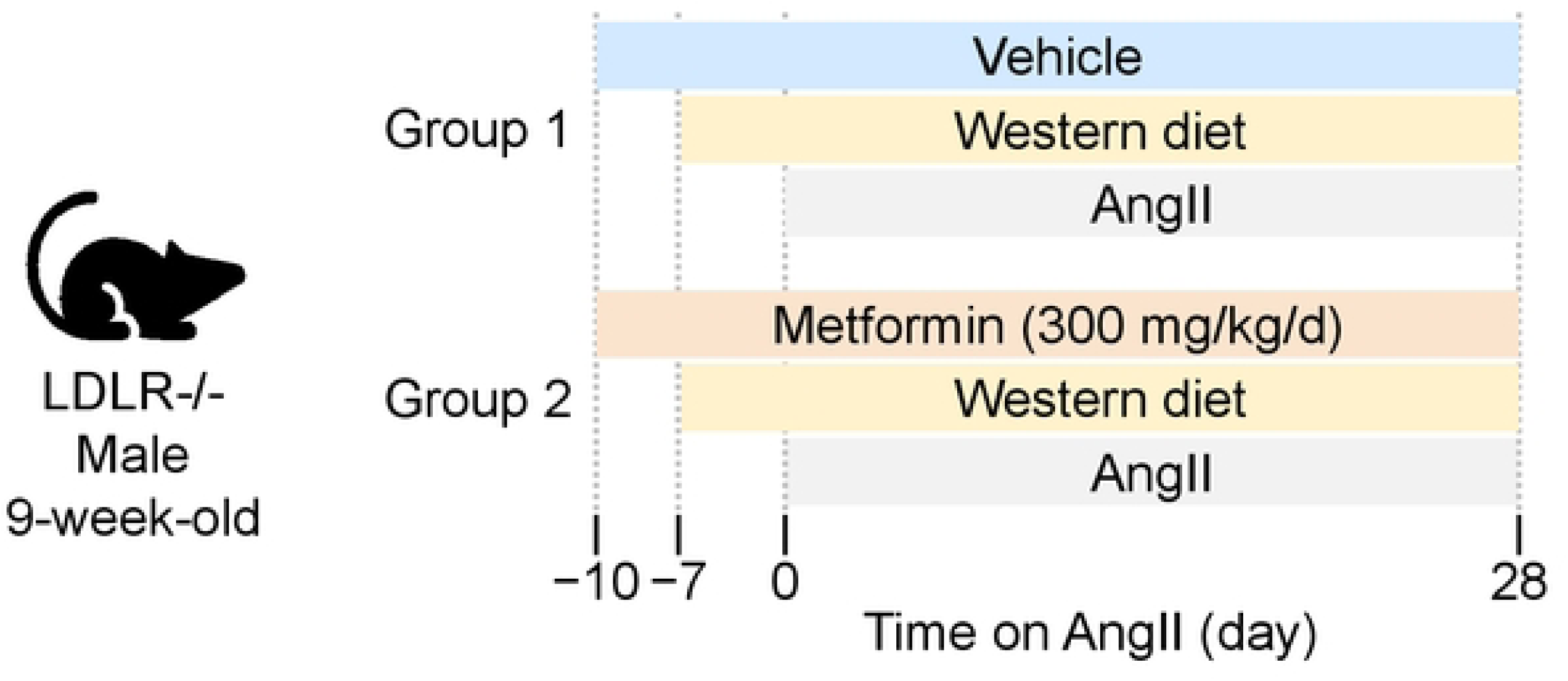
Study design. Angiotensin II (AngII) was infused subcutaneously for 28 days to male low-density lipoprotein receptor-deficient (LDLR−/−) mice administered either vehicle or metformin in drinking water.

### Measurement of Serum Metformin Concentrations

Metformin was extracted from serum via liquid-liquid extraction. Extraction was performed by adding of formic acid (0.1% wt/vol; 980 µL) in acetonitrile to each sample. The samples were then vortexed for 3 minutes and centrifuged at approximately 10,000 rpm for 15 minutes. The supernatant was transferred to a clean glass test tube and evaporated to dryness at 40°C under a stream of nitrogen. The extracted analyte was reconstituted in 100 µL of reconstitution solution, vortexed, and transferred to a silanized glass insert in a labeled vial. Chromatographic separation was performed on a Waters XBridge C18 column (3.0 mm x 50 mm, 3.5 µm particle size). The mobile phase consisted of ammonium acetate (2 mM) in formic acid (0.1% wt/vol) in water as mobile phase A and acetonitrile (100% volvol) for mobile phase B. Compounds were separated across a linear gradient from 1% mobile phase B to 100% mobile phase B over 5 minutes. The flow rate was set to 0.35 ml/min. The column temperature was set to 40°C. Detection was performed via multiple reaction monitoring in positive electrospray ionization mode.

### Quantification of aortic aneurysm

Mice were euthanized by intraperitoneal injection of overdose ketamine (90 mg/kg, #11695-6840-1, Covetrus) and xylazine (10 mg/kg, #11695-4024-1, Covetrus) mixture. After exsanguination via cardiac puncture, the right atrium of the heart was cut to allow perfusate drainage. Saline (8-10 mL) was perfused via the left ventricle to remove blood from the aorta. Then, the aorta was dissected out carefully and placed in neutral buffered formalin (10% wt/vol) for at least 24 hours. Subsequently, perivascular tissues were removed gently from aortas using fine forceps. Entire aortas were then pinned on a black rubber plate, photographed using a digital camera (DS-Ri1, Nikon), and maximum aortic diameters were measured in the ascending and supra-renal abdominal regions using NIS Elements AR 4.51 software (Nikon, Japan). Aortic samples were subsequently cut longitudinally to expose the intimal surface to capture en face images using the digital camera. Ascending aortic areas were traced manually on the en face images, as described previously.[22] Measurements were verified by an individual who was blinded to the identity of the study groups.

### Statistics

Data are presented as individual data points with the median and 25th/75th percentiles. Normality and homogeneity of variance were assessed using the Shapiro-Wilk and Brown-Forsythe tests, respectively. Welch’s t-test or Mann-Whitney U test was used for parametric and non-parametric comparisons between two groups. Since the data met the assumptions of normality and homogeneous variance, a two-way ANOVA followed by the Holm-Sidak test was applied for comparisons among four groups. The incidence of aortic rupture was tested by Fisher’s exact test. P value < 0.05 was considered statistically significant. Statistical analyses were performed using SigmaPlot version 15.0 (SYSTAT Software Inc.).

### Data availability

All numerical data are provided in the Supplemental Excel File. Raw mass spectrometry data are available from the corresponding author.

## RESULTS

### Determination of metformin delivery in AngII-infused mice

We first measured plasma metformin concentrations by mass spectrometry analysis to determine plasma concentrations of AngII-infused mice (**Figure 2A**). As expected, metformin was undetectable in plasma of vehicle-administered mice, whereas metformin-administered mice exhibited concentrations ranging from 95 to 217 ng/mL. In addition, metformin-administered mice showed a significantly lower body weight after 4 weeks of AngII infusion compared to vehicle-administered mice (**Figure 2B**). These findings confirm the effective delivery of metformin in AngII-infused mice.

**Fig 2.**
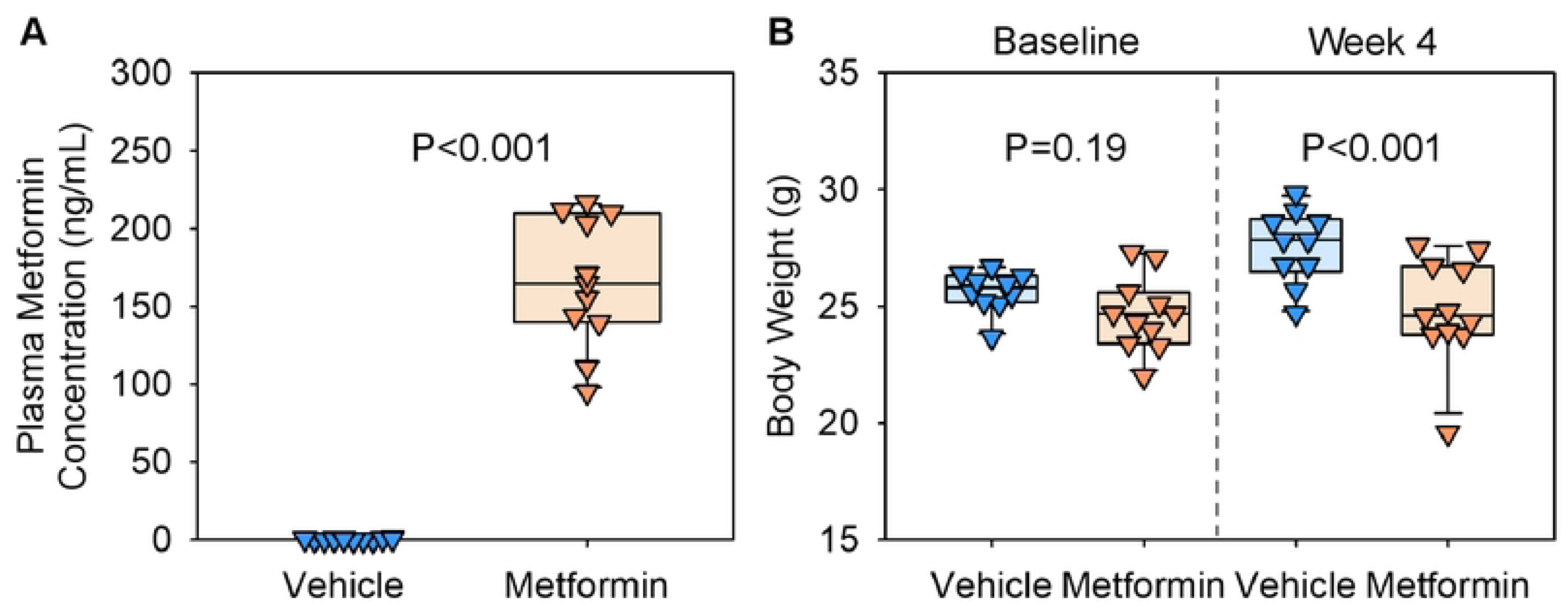
Verification of metformin administration in AngII-infused mice. **(A)** Plasma metformin concentrations measured by mass spectrometry and **(B)** body weight of AngII-infused mice administered either vehicle or metformin in drinking water. Body weight was measured at the baseline and after 4 weeks of AngII infusion. N=10, 11/group. P-values were determined by Welch’s t-test (A) and two-way ANOVA followed by Holm-Sidak test (B).

### Metformin failed to attenuate AngII-induced aortic aneurysm in mice

During the 4-week AngII infusion period, two vehicle-administered mice and one metformin-administered mouse died from suprarenal abdominal aortic rupture. The incidence of aortic rupture did not differ between groups (P = 0.99, Fisher’s exact test). Mice that survived the 28-day infusion were euthanized, and abdominal aortic diameters were measured ex vivo (**Figure 3A**). No significant differences in maximal aortic diameters were observed between vehicle- and metformin-administered mice (**Figure 3B**). Since AngII infusion induces aneurysm formation in the ascending aorta, we also assessed the impact of metformin in this region.[23] Ex vivo measurements revealed no differences in ascending aortic diameters between groups (**Figure 4A**). These findings were further supported by intimal area measurements of the ascending aortic region, which showed comparable ascending aortic areas between the groups (**Figure 4B**). Collectively, these data indicate that metformin did not inhibit AngII-induced aortic aneurysm formation in both the suprarenal and ascending aortic regions.

**Fig 3.**
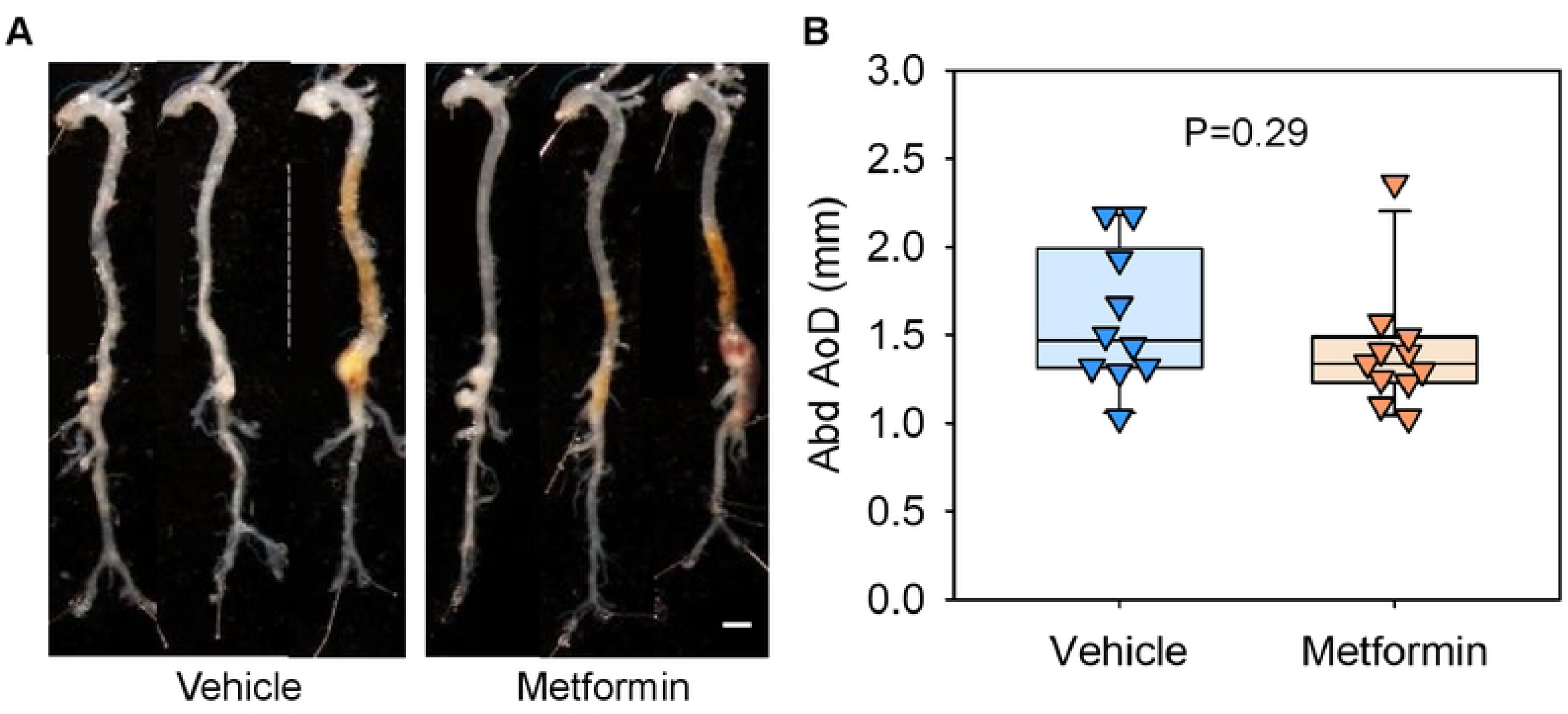
Metformin did not suppress AngII-induced abdominal aortic aneurysms in LDLR−/− mice. **(A)** Representative ex vivo images of aortas and ex vivo measurements of **(B)** abdominal (Abd) aortic diameters (AoD) in AngII-infused LDLR−/− mice fed a Western diet with either vehicle or metformin administration. Scale bar = 2 mm. P-values were determined by Welch’s t-test

**Fig 4.**
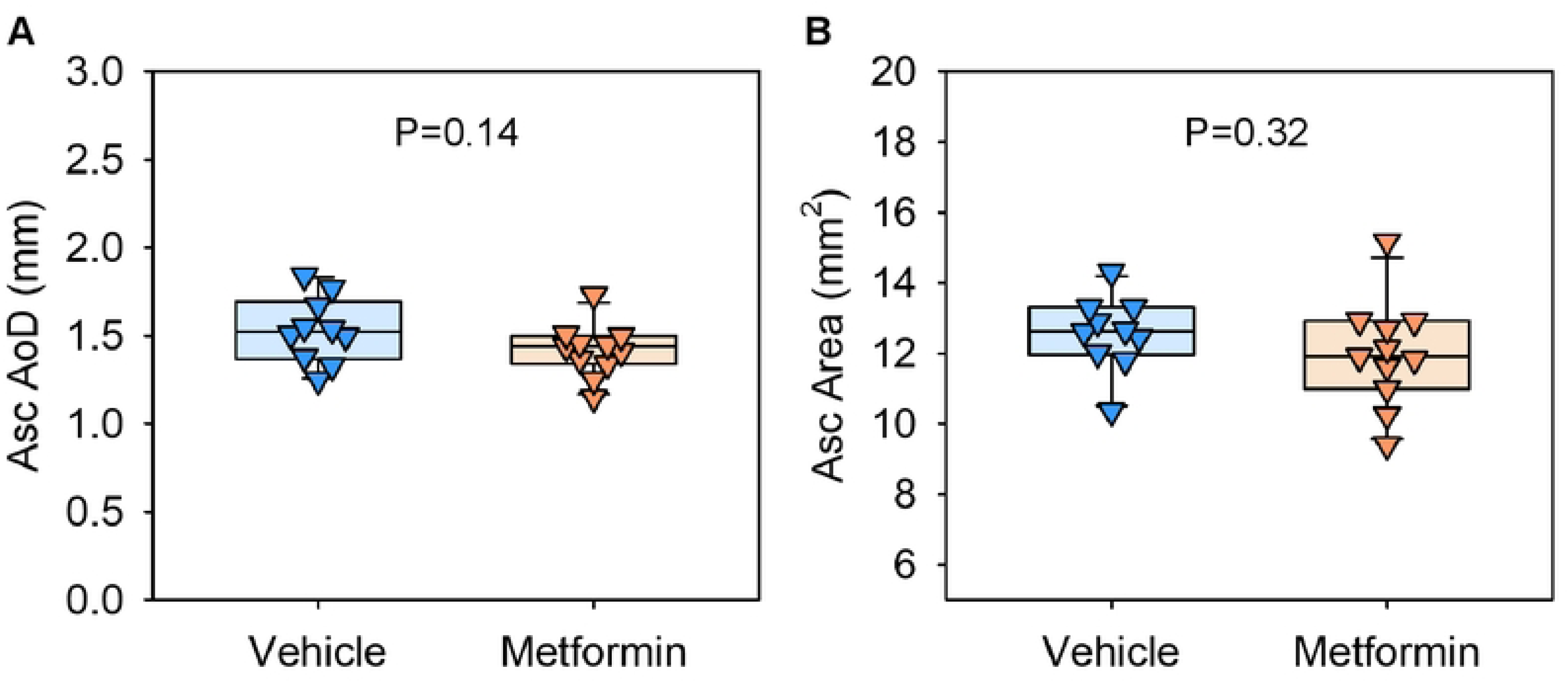
Metformin did not suppress AngII-induced thoracic aortic aneurysms in LDLR−/− mice. Ex vivo measurements of ascending (Asc) aortic **(A)** diameters (AoD) and **(B)** areas in AngII-infused LDLR−/− mice fed a Western diet with either vehicle or metformin administration. Scale bar = 2 mm. P-values were determined by Welch’s t-test.

## DISCUSSION

While metformin has been shown to reduce AngII-induced AAA formation in ApoE−/− mice in previous studies,[13, 14] it failed to exert similar protective effects in LDLR−/− mice in the present study. Both ApoE−/− and LDLR−/− mice develop hypercholesterolemia; however, their lipoprotein profiles differ significantly.[24] ApoE−/− mice exhibit markedly elevated cholesterol concentrations carried in VLDL and chylomicron remnants, with minimal HDL cholesterol concentrations, even when fed normal laboratory diet.[19] In contrast, LDLR−/− mice develop hypercholesterolemia characterized by increased LDL cholesterol concentrations when fed a high-fat or Western diet.[19] These differences may influence the pathophysiology of AAAs, resulting in the alteration of AAA susceptibility in ApoE−/− mice. Another important factor is glucose metabolism. Diabetogenic diet increases plasma glucose concentrations in LDLR−/− mice, whereas ApoE−/− mice are resistant to diet-induced hyperglycemia.[20] Differences in glucose homeostasis between these models may contribute to the divergent outcomes of metformin. Further studies are required to investigate the specific contributions of glucose metabolism in AAA formation and the precise conditions under which metformin plays its vascular protective effects.

In this study, metformin was administered in drinking water. Based on the standard water intake of mice (6 mL water/day),[25] mice were predicted to receive approximately 300 mg/kg of metformin per day. This dose is comparable to or higher than those used in previous studies demonstrating protective effects of metformin against AAA in mice (100 and 250 mg/kg).[26] In the present study, effective drug delivery was determined by measuring plasma metformin concentrations and body weight. Mass spectrometry validated the presence of metformin in the plasma of metformin-administered mice compared to controls. These results are direct evidence showing drug exposure.

Metformin-administered mice had significantly lower body weight than vehicle-administered mice. Given that metformin decreases body weight in mice,[27] these data suggest that metformin was effectively delivered and had physiological effects. Nonetheless, whether the administered dose was sufficient to influence AngII-induced TAA and AAA formation remains an open question.

Retrospective clinical studies have shown that metformin use is associated with reduced prevalence, growth, and rupture of AAAs in patients with diabetes.[12, 28, 29] However, the mechanistic basis for this association remains unclear. Investigating the impact of diabetes on AAA pathophysiology is challenging due to multiple confounding factors. The mechanisms by which hyperglycemia, elevated plasma insulin concentrations, and individual anti-diabetic medications influence AAA pathophysiology remain poorly understood. Our findings that metformin does not suppress AngII-induced AAA in LDLR−/− mice contrast with the retrospective clinical studies indicating protective effects of metformin in AAAs. These highlight the need for further research to elucidate the complex relationship between diabetes and AAA pathophysiology.

In conclusion, the present study revealed that metformin did not attenuate either AngII-induced ascending or abdominal aortic aneurysms in LDLR−/− mice.

## Acknowledgments

Plasma metformin concentrations were measured by mass spectrometry in the Mass Spectrometry and Proteomics Core at the University of Kentucky (RRID:SCR_026359)

## AUTHOR DECLARATIONS

## Authors’ contributions

SCT and HS analyzed the data, wrote the main manuscript text, and prepared figures. SCT, SI, JCH, MKF, and DAH implemented animal experiments. SCT, HSL, AD, and HS contributed to designing studies. All authors reviewed the manuscript.

